# Unravelling Representations in Scene-selective Brain Regions Using Scene Parsing Deep Neural Networks

**DOI:** 10.1101/2020.03.10.985309

**Authors:** Kshitij Dwivedi, Radoslaw Martin Cichy, Gemma Roig

**Author notes:** jointly directed work (Radoslaw Martin Cichy, Gemma Roig).

## Abstract

Visual scene perception is mediated by a set of cortical regions that respond preferentially to images of scenes, including the occipital place area (OPA) and parahippocampal place area (PPA). However, the differential contribution of OPA and PPA to scene perception remains an open research question. In this study, we take a deep neural network (DNN)-based computational approach to investigate the differences in OPA and PPA function. In a first step we search for a computational model that predicts fMRI responses to scenes in OPA and PPA well. We find that DNNs trained to predict scene components (e.g., wall, ceiling, floor) explain higher variance uniquely in OPA and PPA than a DNN trained to predict scene category (e.g., bathroom, kitchen, office). This result is robust across several DNN architectures. On this basis, we then determine whether particular scene components predicted by DNNs differentially account for unique variance in OPA and PPA. We find that variance in OPA responses uniquely explained by the navigation-related floor component is higher compared to the variance explained by the wall and ceiling components. In contrast, PPA responses are better explained by the combination of wall and floor, that is scene components that together contain the structure and texture of the scene. This differential sensitivity to scene components suggests differential functions of OPA and PPA in scene processing. Moreover, our results further highlight the potential of the proposed computational approach as a general tool in the investigation of the neural basis of human scene perception.

## 1 Introduction

Visual scene understanding is a fundamental cognitive ability that enables humans to interact with the components and objects present within the scene. Within the blink of an eye we know what type of scene we are in (e.g. kitchen, or outdoors), as well as its spatial layout and the objects contained in it.

Research on the neural basis of scene understanding has revealed a set of cortical regions with a preferential response to images of scenes over images of objects. These regions are the parahippocampal place area (PPA) [Epstein and Kanwisher, 1998], occipital place area (OPA) [Dilks et al., 2013, Hasson et al., 2003], and retrosplenial cortex (RSC) [O’Craven and Kanwisher, 2000]. To investigate the distinct function each of these place regions has, subsequent research has begun to tease apart their commonalities and differences in activation profile and representational content [Epstein and Kanwisher, 1998, Hasson et al., 2003, Dilks et al., 2013, Bonner and Epstein, 2017, Silson et al., 2015, O’Craven and Kanwisher, 2000]. However, a complete picture of how scene-selective regions together orchestrate visual scene understanding is still missing.

To gain further insights, a promising, but relatively less explored approach is computational modelling of brain activity. Recently, large advances have been made in modeling activity in visual cortex using deep neural networks (DNNs) trained on object categorization tasks [Krizhevsky et al., 2012] in both human and non-human primates [Yamins et al., 2014, Khaligh-Razavi and Kriegeskorte, 2014, Cichy et al., 2016]. Inspired by this success, researchers have also begun to use DNNs trained on scene categorization to investigate scene-selective cortex [Cichy et al., 2017, Bonner and Epstein, 2018, Groen et al., 2018].

In this process two issues have emerged that need to be addressed. First, while DNN trained on categorization tasks currently do best in predicting activity in scene-selective cortical regions, they do not account for all explainable variance. One particularly promising direction is the exploration of models trained on tasks different from categorization that might more closely resemble the brain region’s functionality, and thus predict brain activity better [Yamins et al., 2014, Cichy and Kaiser, 2019]. Second, it remains unclear what is the nature of the representations in the DNNs that gives them their predictive power. Thus, additional effort is needed to clarify what these representations are.

To address the above issues, we investigated neural activity in the scene-selective cortex using DNNs trained on scene parsing instead of categorization. A scene parsing task requires the DNN to predict the location and category of each scene component in the image. While the scene-categorization task requires only recognizing the scene category, the scene parsing task requires deeper scene understanding involving categorization as well as a grasp of the spatial organization of components and objects within the scene. In order to help interact with different objects and navigate within the scene, scene-selective brain regions should also encode the spatial organization of components within the scene. Therefore, we hypothesize that the scene parsing task is closer to the task the brain has to solve, and a DNN trained on scene parsing will predict brain activity better than a DNN trained on scene categorization.

To evaluate our hypothesis, we compared the power of DNNs trained on scene parsing versus categorization to predict activity in scene-selective cortical regions. For this we used an existing fMRI set of brain responses elicited by viewing scene images [Bonner and Epstein, 2017] and applied representational similarity analysis (RSA) to compare brain responses with DNNs. We found that scene parsing DNNs explain significantly more variance in brain responses uniquely in scene-selective regions than scene-classification DNNs.

We next investigated what representations in the DNNs trained on scene parsing gave the model its predictive power. For this we queried the DNN’s representations of different scene components, considering components that were present in all stimulus images: wall, floor, and ceiling. We showed that different scene components predict responses in OPA and PPA differently: floor explained more variance in OPA than wall and ceiling, while wall explained more variance in PPA than floor and ceiling. Importantly, results were consistent across three different DNN architectures, showing the generalizability of our claims across architectures.

In sum, our results reveal differential representational content in scene-selective regions OPA and PPA, and highlight DNNs trained on scene parsing as a promising model class for modelling human visual cortex with well interpretable output.

## 2 Materials and Methods

### 2.1 fMRI data

We used fMRI data from a previously published study by Bonner and Epstein [2017] where all experimental details can be found, as well as instructions on how to download the data. The fMRI data were collected from 16 participants on a Siemens 3.0T Prisma scanner with a 64-channel head coil. The participants were presented with images of indoor environments and performed a category-detection task. The images were presented for 1.5s on the screen followed by a 2.5s interstimulus interval. The images presented in the experiment were from a stimulus set of 50 color images depicting indoor environments. Voxel-wise (voxel size = 2 × 2 × 2 mm) responses to each image during each scan run were extracted using a standard linear model.

We here focus on two scene-selective regions of interest (ROIs): PPA and OPA. PPA and OPA were identified from separate functional localizer scans using a contrast of brain responses to scenes larger than to objects and additional anatomical constraints. For both ROIs and all the subjects, each voxel’s responses in a given ROI were z-scored across images in a given run and then averaged across runs. The responses to a particular image were further z-scored across voxels.

### 2.2 Behavioral data

We used scene-related behavioral data representing navigational affordances assessed on the same stimulus set as used for recording the fMRI data described above [Bonner and Epstein, 2017]. To represent navigational affordances, a behavioral experiment was conducted in which 11 participants (different from the participants in the fMRI experiment) indicated the path to walk through the indoor environment used in the fMRI study using a computer mouse. The probabilistic maps of paths for each image were created, followed by a histogram construction of navigational probability in one-degree angular bins radiating from the bottom center of the image. These histograms represent a probabilistic map of potential navigation routes from the viewer’s perspective. The resultant histogram is referred to as the Navigational Affordance Model (NAM).

### 2.3 DNN Models

We selected DNNs optimized on two different scene-related tasks: scene classification and scene parsing. We describe both types of models in detail below.

#### Scene-classification models

For solving a scene-classification task, a DNN model is optimized to predict the probabilities of the input image belonging to a particular scene-category. For comparison with neural and behavioral data, we considered DNNs pretrained on the scene-classification task on the Places-365 dataset [Zhou et al., 2017]. Places-365 is a large scale scene-classification dataset consisting of 1.8 million training images from 365 scene categories. We selected multiple scene-classification DNN architectures to investigate if our results generalize across different architectures. For this purpose, we considered 3 standard architectures: Alexnet [Krizhevsky et al., 2012], Resnet-18 [He et al., 2016], and Resnet-50 [He et al., 2016] and downloaded pretrained models from https://github.com/CSAILVision/places365.

Alexnet consists of 5 convolutional layers (conv1-conv5) followed by 3 fully connected layers (fc6, fc7, and fc8). Both Resnet-18 and Resnet-50 consist of a convolutional layer followed by four residual blocks (block1 - block4) each consisting of several convolutional layers with skip connections leading to a final classification layer (fc). Resnet-18 consists of 18 layers and Resnet-50 consists of 50 layers in total and they differ in the number of layers within each block.

#### Scene parsing models

We used scene parsing models trained on ADE20k scene parsing dataset [Zhou et al., 2016]. The ADE20k dataset is a densely annotated dataset consisting of 25k images of complex everyday scenes with pixel-level annotations of objects and components.

For the first set of experiments, where we compare the predictive power of scene parsing models to scene-classification models for explaining the neuronal responses, we design scene parsing models such that their encoder architecture is taken from scene-classification models while their decoder architecture is task-specific. The encoder of the scene parsing models consists of the convolutional part (conv1-conv5 of Alexnet, and block1-block4 of Resnet18 and Resnet50) of scene classification models. The decoder of scene parsing models is adapted to the scene parsing task following the architecture proposed by Zhao et al. [2017]. It consists of a Pyramid Pooling module with deep supervision [Zhao et al., 2017] (d1), followed by a layer (d2) that predicts several spatial maps, one spatial map per scene component predicted, that represent the probability of the presence of that component at a given spatial location. The encoder weights of scene parsing models are initialized with the weights learned on the scene-classification task and decoder weights are initialized randomly. The scene parsing DNNs are then trained on ADE20k training data using a per-pixel cross-entropy loss. The above procedure ensures that gain/drop in explaining neural/behavioral responses could only be due to additional supervision on the scene parsing task.

The aforementioned scene parsing DNNs are well suited for a direct comparison with the scene categorization DNNs as they have the same encoder architecture and were initialized with weights learned on scene-categorization task. However, they are not comparable to state-of-the-art models in terms of accuracy on the scene parsing task. Since our aim is to reveal differences in representations of the scene areas in the brain by comparing scene components, for the second set of experiments, it is crucial to select components detected with DNNs from the literature that achieve the highest accuracy in scene parsing. For this reason, we selected 3 state-of-the-art models on the scene parsing task namely Resnet101-PPM [Zhou et al., 2016], UperNet101 [Xiao et al., 2018], and HR-Netv2 [Sun et al., 2019]. We selected multiple models to investigate if the results we obtain are consistent across different models. Resnet101-PPM consists of a dilated version of the Resnet101 model (a deeper version of Resnet50 that consists of a total of 101 layers) trained on Imagenet as the encoder and a Pyramid Pooling module with deep supervision [Zhao et al., 2017] as the decoder. Upernet101 [Xiao et al., 2018] is based on the Feature Pyramid Network by Lin et al. [2017] that uses multi-level feature representations via a top-down architecture to fuse high-level semantic features with mid and low-level using lateral connections. Upernet101 also has a Pyramid Pooling Module before the top-down architecture to overcome the small receptive-field issue. HRNetv2 [Sun et al., 2019] relies on the importance of high-resolution feature maps for pixel labeling maps by maintaining high-resolution feature representations throughout the architecture by high-to-low resolution convolutions in parallel. We downloaded all above mentioned models from https://github.com/CSAILVision/semantic-segmentation-pytorch.

To reveal performance differences between different models on the scene parsing task, we compared the performance of state-of-the-art models with the scene parsing models used for comparison with scene-classification DNNs (see above, Alexnet, Resnet-18, Resnet-50). For the comparison, we calculated the mean intersection over union (mIoU) score of detecting all components for all the images from the ADE20k validation dataset. The IoU score is calculated by dividing the intersection between a predicted and corresponding ground truth pixel-level segmentation mask by their union. IoU is a standard metric to evaluate the overlap of a predicted and corresponding pixel-level mask of a particular component. Mean IoU is calculated by taking the mean of IoU scores across all images in the validation dataset for all components.

As illustrated in Figure 1a, a scene parsing model decomposes an image into its constituent components. This decomposition allows investigating which scene components are more relevant to explaining the representations in scene-selective brain regions. We first identified which scene components are present in all the images from the stimulus set used for obtaining fMRI responses. To achieve this, we feedforwarded all the 50 images in the stimulus set of the fMRI dataset through the models and checked the presence of all the components in the image. Since the DNN has been trained on an image dataset that is different from the set of stimuli used for the fMRI data, not all scene components predicted by the DNN appear in the stimulus set. In this particular set, we found that wall, floor, and ceiling were core scene components present in all images.

**Figure 1:**
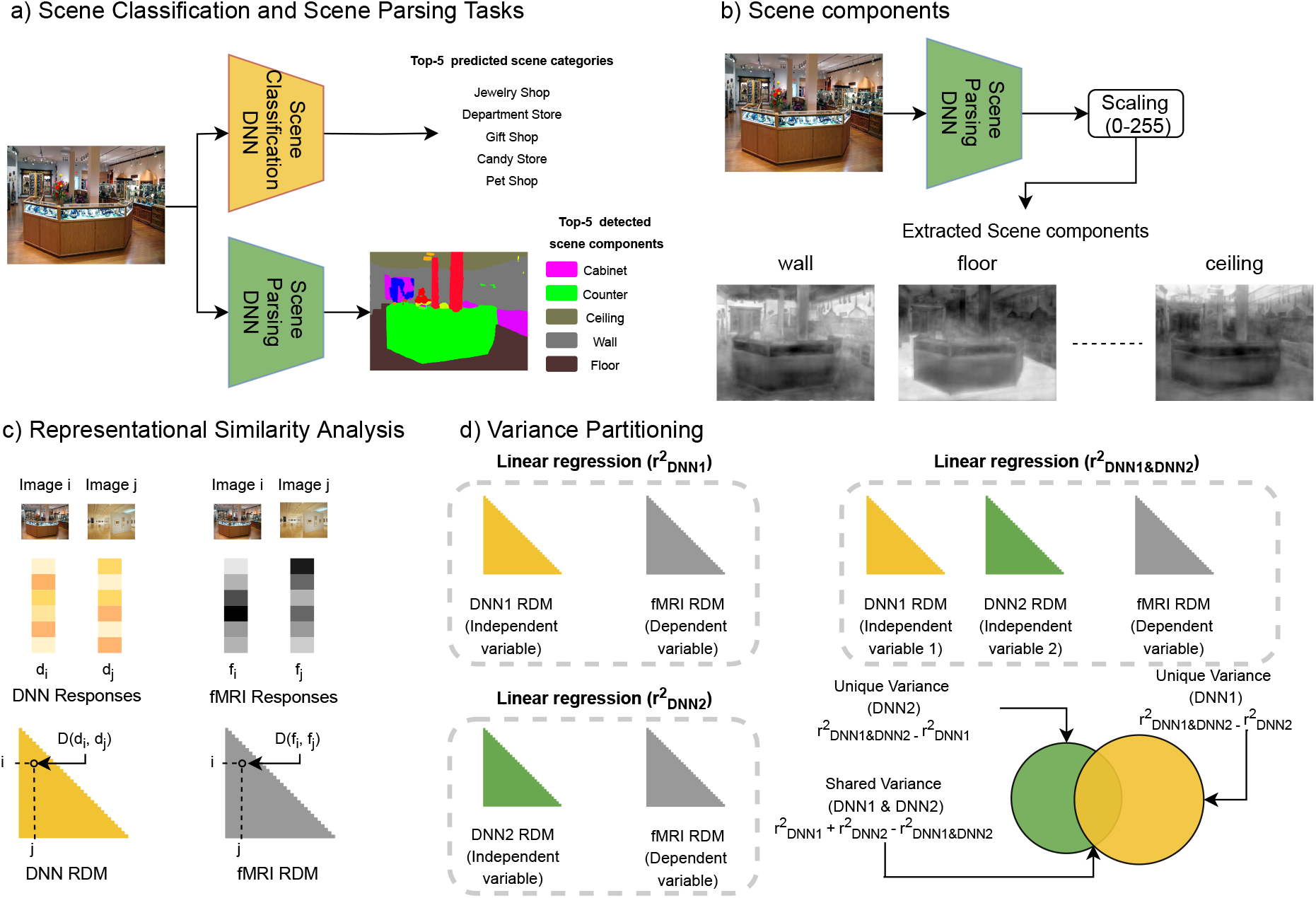
Outline of our approach. a) In the scene classification task, the model outputs the probability of an image belonging to a particular class. In the scene parsing task, the model outputs a spatial map for each component. The pixel value of the spatial map corresponding to a component represents the probability of that pixel belonging to that component. b) We use DNNs trained on scene parsing to extract responses corresponding to individual scene components. c) RSA: We first compute RDMs for a DNN model and a brain ROI by computing pairwise distance (D) between DNN (d_i_,d_j_)/fMRI(f_i_,f_j_) responses corresponding to each pair (i,j) of images in the stimulus set. We next compute the correlation of a DNN RDM with fMRI RDM to determine the similarity between the brain and the DNN. d) Variance Partitioning: We conduct three multiple linear regressions with DNN RDMs as the independent variables and fMRI RDM as the dependent variable to estimate unique and shared variance of fMRI RDM explained by DNN RDMs.

A scene parsing model outputs a spatial probability map for each component. To scale the spatial probability maps corresponding to different components in the same range, we normalized the spatial probability map for each component independently such that each pixel value lies in the range [0, 255]. We show the extracted normalized scene components corresponding to the wall, floor, and ceiling components for an example stimulus in Figure 1b.

### 2.4 Representational Similarity Analysis (RSA)

We applied representational similarity analysis (RSA; [Kriegeskorte et al., 2008] to compare DNN activations and scene components with neural and behavioral responses. RSA enables relating signals from different source spaces (such as here behavior, neural responses, DNN activation) by abstracting signals from separate source spaces into a common similarity space. For this, in each source space condition-specific responses are compared to each other for dissimilarity (e.g., by calculating Euclidian distances between signals) and the values are aggregated in so-called representational dissimilarity matrices (RDMs) indexed in rows and columns by the conditions compared. RDMs thus summarize the representational geometry of the source space signals. Different from source space signals themselves RDMs from different sources spaces are directly comparable to each other for similarity and thus can relate signals from different spaces. We describe the construction of RDMs for different modalities and the procedure by which they were compared in detail below.

#### fMRI ROI RDMs

First, for each ROI (OPA, and PPA), individual subject RDMs were constructed using Euclidean distances between the voxel response patterns for all pairwise comparisons of images. Then, subject-averaged RDMs were constructed by calculating the mean across all individual subject RDMs. We downloaded the subject averaged RDMs of OPA and PPA from the link (https://figshare.com/s/5ff0a04c2872e1e1f416) provided in Bonner and Epstein [2018].

#### Navigational affordance model (NAM) RDMs

NAM RDMs were constructed using Euclidean distances between the navigational affordance histograms for all pairwise comparisons of images. We downloaded the NAM RDM from (https://figshare.com/s/5ff0a04c2872e1e1f416).

#### DNN RDMs

For all the DNNs we investigated in this work, we constructed the RDM for a particular layer using 1−*ρ*, where *ρ* is the Pearson’s correlation coefficient, as the distance between layer activations for all pairwise comparisons of images. For scene classification DNN RDMs, we created one RDM for each of the 5 convolutional layers (conv1-conv5) and for the 3 fully connected layers (fc6,fc7, and fc8) for Alexnet, and the last layer of each block (block1 - block4) and the final classification layer (fc) of Resnet-18/Resnet-50 to compare with neural/behavioral RDMs. For scene parsing DNN RDMs, we created one RDM for each of the 5 convolutional layers (conv1-conv5) and for the 2 decoder layers (d1 and d2) for Alexnet, and the last layer of each block (block1 - block4) and 2 decoder layers (d1 and d2) of Resnet-18/Resnet-50 to compare with neural/behavioral RDMs.

#### Scene component RDMs

For each of the scene components investigated we constructed RDM for it using 1−*ρ* as the distance between normalized spatial probability maps of that scene component, based on all pairwise comparisons of images.

#### Comparing DNN and scene component RDMs with behavioral and neural RDMs

In this work, we pose two questions: first, whether scene parsing models can better explain scene-selective neural responses and navigational affordance behavioral responses better than scene-classification models, and second, whether the scene-components detected by scene parsing models reveal differences in representations of scene-selective ROIs.

To investigate the first question, we calculated the Spearman’s correlation between the RDMs of different layers of a scene-classification DNN with a particular behavioral/neural RDM and selected the layer RDM that showed the highest correlation with the behavioral/neural RDM. We used the selected layer RDM as the representative RDM for that architecture. We repeated the same procedure to select the representative RDM from a scene parsing model. We report the layer used to select the representative RDMs for each model in Table 1. To compare which model RDM (scene parsing or scene-classification) explains behavioral/neural RDM better, we compared the correlation values of both RDMs with behavioral/neural RDM (Figure 1c).

**Table 1:** Correlation value and layer information of the layer that showed the highest correlation with a particular brain area or behavior for all the models considered (AlexNet, ResNet18 and ResNet50).

| Task | Models | OPA | PPA | NAM |
| --- | --- | --- | --- | --- |
| Scene class. | Alexnet | 0.395(fc7) | 0.369(fc7) | 0.122(conv5) |
|  | Resnet18 | 0.391(fc) | 0.397(fc) | 0.114(block4) |
|  | Resnet50 | 0.364(fc) | 0.365(fc) | 0.111(block4) |
| Scene parsing | Alexnet | 0.393(d2) | 0.393(d1) | 0.162(conv5) |
|  | Resnet18 | 0.426(d1) | 0.415(d1) | 0.198(block4) |
|  | Resnet50 | 0.415(d1) | 0.418(d1) | 0.165(d2) |

To investigate whether the scene-components detected by scene parsing models reveal differences in representations of scene-selective ROIs, we computed the correlation between a scene component RDM and a neural RDM and compared which scene component explains better a particular ROI.

### 2.5 Variance Partitioning

While in its basic formulation RSA provides insights about the degree of association between a DNN RDM and a behavioral/neural RDM, it does not provide a full picture of how multiple DNN RDMs together explain the behavioral/neural RDM. Therefore, we applied a variance partitioning analysis that determines the unique and shared contribution of individual DNN RDMs in explaining the behavioral/neural RDM when considered in conjunction with the other DNN RDMs.

We illustrate the variance partitioning analysis in Figure 1d. We assigned a behavioral/neural RDM as the dependent variable (referred to as predictand). We then assigned two model (DNN/scene component) RDMs as the independent variables (referred to as predictors). Then, we performed three multiple regression analyses: one with both independent variables as predictors, and two with individual independent variables as the predictors. Then, by comparing the explained variance (r^2^) of a model used alone with the explained variance when it was used with other models, the amount of unique and shared variance between different predictors can be inferred (Figure 1d).

To compare scene parsing and scene-classification models, the predictors were the respective DNN RDMs and predictands were the behavioral and neural RDMs. To compare different scene components, the predictors were the respective scene component RDMs and predictands were the neural RDMs of scene-selective ROIs.

### 2.6 Statistical Testing

We applied nonparametric statistical tests to assess the statistical significance in a similar manner to a previous related study [Bonner and Epstein, 2018]. We assessed the significance of the correlation between neural/behavioral responses with a DNN through a permutation test by permuting the conditions randomly 5000 times in either the neural ROI RDM or the DNN RDM. From the distribution obtained using these permutations, we calculated p-values as one-sided percentiles. We calculated the standard errors of these correlations by randomly resampling the conditions in the RDMs for 5000 iterations. We used re-sampling without replacement by subsampling 90% (45 out of 50 conditions) of the conditions in the RDMs. We used an equivalent procedure for testing the statistical significance of the correlation difference and unique variance difference between the two models.

## 3 Results

### 3.1 Are scene parsing models suitable to account for scene-selective brain responses and scene-related behavior?

We investigated the potential of DNNs trained on scene parsing to predict scene-related human brain activity focusing the analysis on scene-selective regions OPA and PPA. To put the result into context we compared the predictive power of DNNs trained on scene parsing to DNNs trained on scene classification, which are currently the default choice in investigating scene-related brain responses and behavior [Bonner and Epstein, 2018, Groen et al., 2018, Cichy et al., 2017]. To ensure that the results can be attributed to differences in the task rather than being specific to particular network architecture, we investigated three different network architectures: Alexnet, Resnet18, and Resnet50.

We applied representational similarity analysis (RSA) to relate DNN models (scene classification and scene parsing) with the brain responses in OPA and PPA (Figure 2a). We found that DNNs trained on scene parsing significantly predicted brain activity in all investigated regions (all p<0.05). This shows that they are suitable candidate models for the investigation of brain function. We further found that DNNs trained on scene parsing explain as much or more variance in scene-selective regions than DNNs trained on scene-categorization across different architectures.

**Figure 2:**
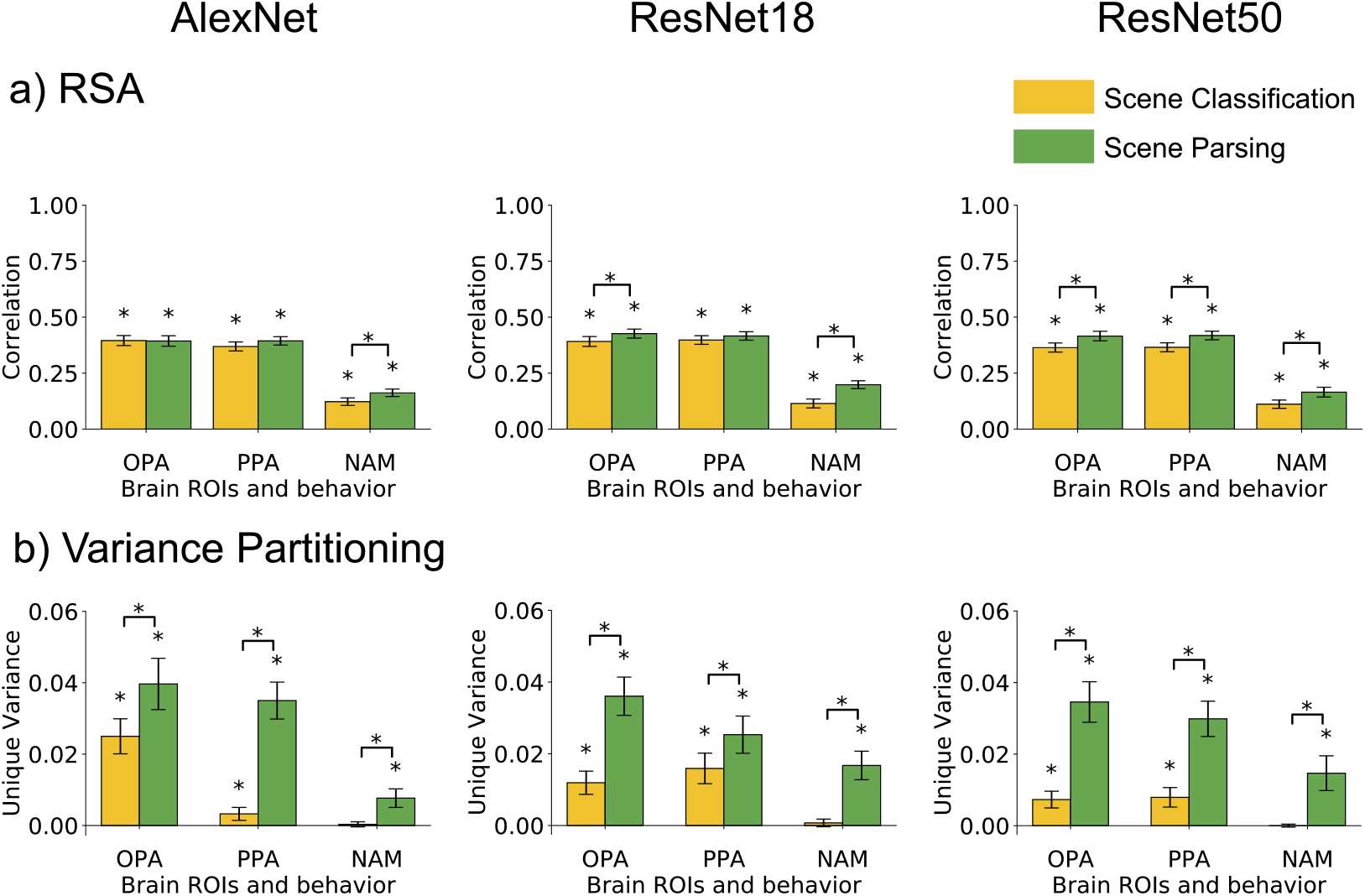
Model comparison in accounting for OPA and PPA as well as behavior. a) RSA of scene-selective areas PPA, OPA, and behavioral model NAM with scene parsing and scene-classification models, and b) variance of scene-selective areas PPA, OPA, and behavioral model NAM explained uniquely by scene parsing and scene-classification models for the architecture Alexnet (left), Resnet18 (middle), Resnet50 (right). The asterisk at the top indicates the significance (* p <0.05) calculated by permuting the conditions 5000 times.

If scene parsing models are suitable models for predicting responses in scene-selective brain regions, and these regions underlie scene understanding, the models should predict scene-related behavior, too. We considered navigational affordance behavior operationalized as the angular histogram of navigational trajectories that participants indicated for the stimulus set. Paralleling the results on brain function, the investigation of behavior showed that DNNs trained on scene parsing predicted behavior significantly (all p<0.05), and also significantly better than DNNs trained on scene-classification (all p<0.05).

While the RSA results above provided insights about the degree of association between a DNN RDM and behavioral/neural RDM, it cannot tell how multiple DNN RDMs together predict the behavioral/neural RDM. For this more complete picture, we conducted variance partitioning to reveal the unique variance of neural/behavioral RDMs explained by scene-classification and scene parsing DNN RDMs (Figure 2b). We observe from Figure 2b that scene parsing DNNs explain more variance uniquely (p<0.05) than scene-classification DNNs for both scene-selective ROIs. We further observe for scene-selective neural RDMs (Figure 3) that most of the variance explained is shared between scene-classification and scene parsing DNNs across all three architectures. The results suggest that scene parsing DNNs might be a better choice for investigating scene-selective neural responses than scene-classification DNNs.

**Figure 3:**
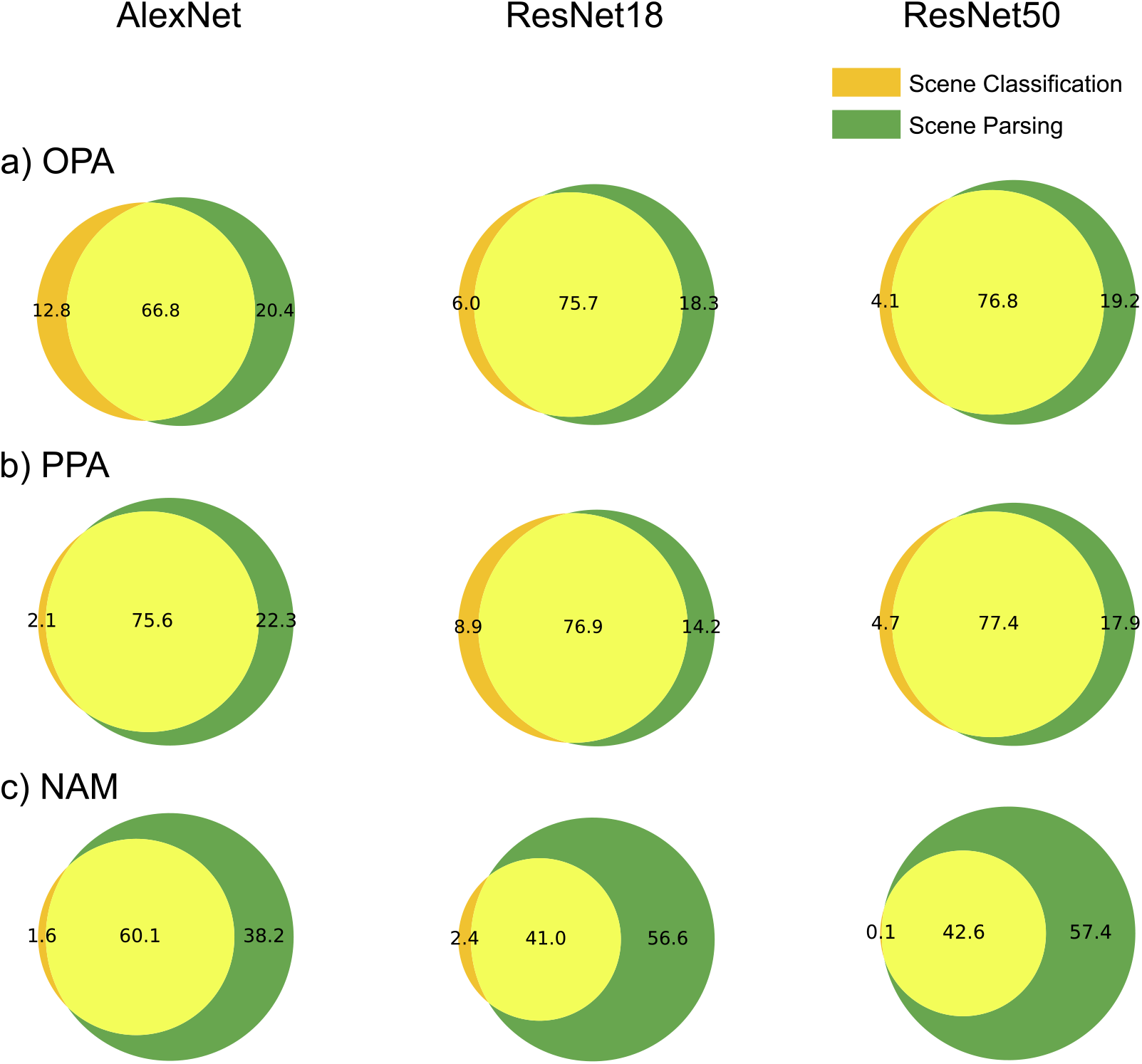
Variance Partition Results of Scene-parsing vs. Scene Classification Comparison. The plots show the unique and shared variance of a) OPA, b) PPA and c) NAM explained by Scene-parsing and Scene-classification DNNs for three architectures: Alexnet (left), Resnet18 (center), and Resnet50 (right).

We observe for behavior that the scene-classification DNNs explain nearly no unique variance, while on the other hand scene parsing DNNs explain significantly higher unique variance (p<0.05) across all three architectures (see Figure 2b for unique variance, and Figure 3 for Venn Diagram illustrating both unique and shared variances). The results suggest that since the scene parsing task takes into account the spatial arrangement of constituent components in the scene, a scene parsing DNN is a better choice for explaining behavioral responses related to the spatial organization of the scene than a scene-classification DNN.

Together, these results establish DNNs trained on scene parsing tasks as a promising model class for investigating scene-selective cortical regions in the human brain and for navigational behavior related to the spatial organization of scene components.

### 3.2 State-of-the-art scene parsing models for investigating scene components represented in the human brain

Models trained on the scene parsing task offer the possibility to selectively investigate which of the scene components (such as the wall, ceiling or floor) they encode. But, what is the most suitable scene parsing model to compare to the brain? In the model comparison above our choice was guided by making models as similar to each other as possible in complexity to rule out that observed differences in accounting for brain activity are simply due to differences in model complexity. However, for in-depth investigation of scene-selective areas using scene-components it is crucial to choose models that detect the scene-components with as high accuracy as possible. Therefore, we compared the performance of different scene parsing models qualitatively and quantitatively to select the most accurate ones to compare with the responses of scene-selective brain areas.

For performance comparison on the scene parsing task, we chose the three models used above (Alexnet, Resnet18, and Resnet 50), plus three state-of-the-art models of scene parsing: HRNetv2, Upernet101, and Resnet101-PPM. The state-of-the-art models achieve high performance by merging low-resolution feature maps with high-resolution feature maps to generate results in high spatial resolution. We illustrate their parsing performance qualitatively by examining their output on an example image (Figure 4). We observe that the scene parsing output generated by Resnet50 had smooth and less precise boundaries of components while Resnet101-PPM, Upernet101, and HRNetv2 detected components accurately with precise boundaries in their outputs.

**Figure 4:**
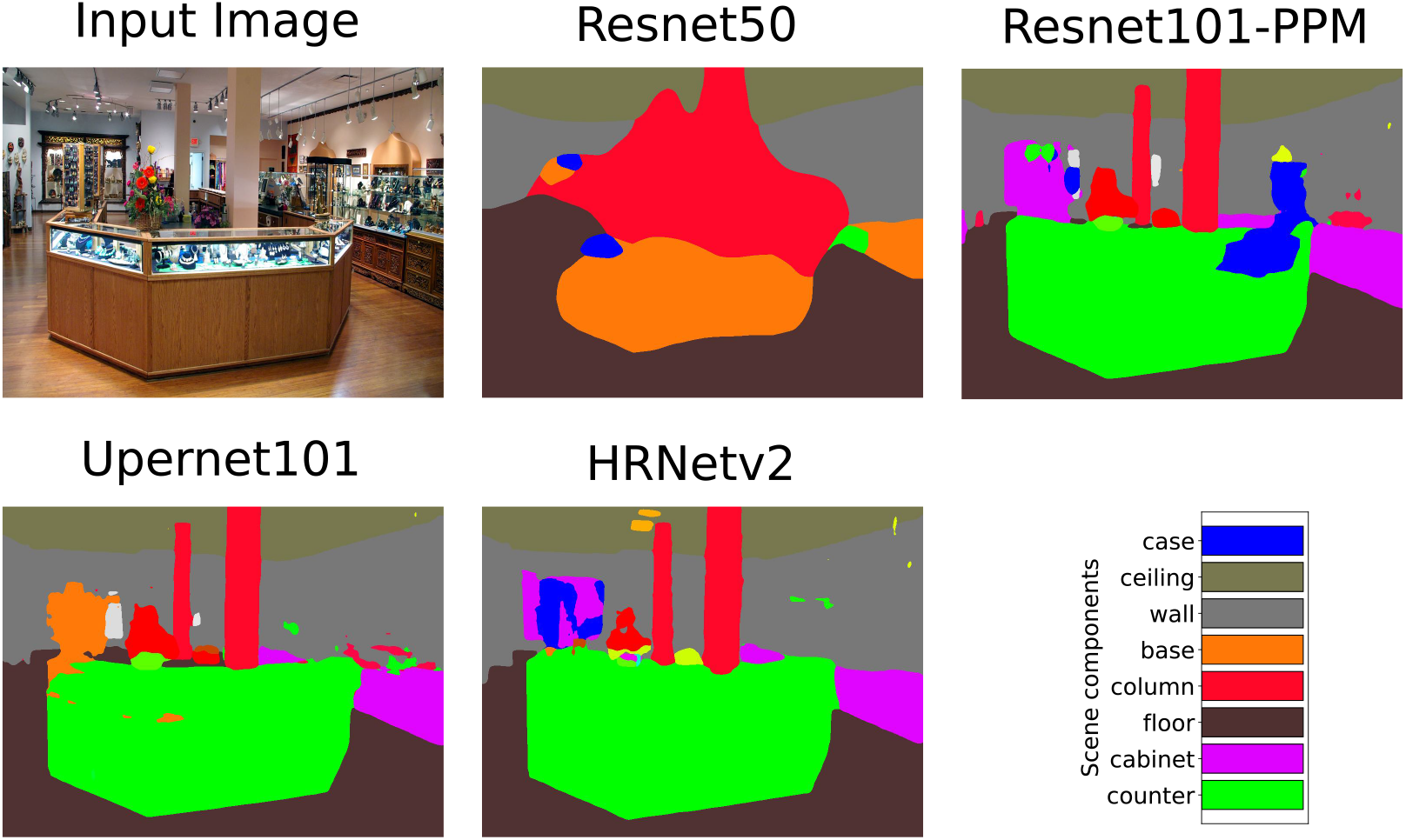
Qualitative comparison of scene parsing output for different models. Input image (top left) and corresponding scene parsing output of the different models investigated in this work.

To quantitatively compare model performance, we evaluated the performance of all models on the ADE20k validation dataset. For this we calculated the mean intersection over union (mIoU) score of detecting all components for all the images from the ADE20k validation dataset. We report mIoU scores of individual components that were present in all images in the stimulus set: wall, ceiling, and floor. The results are reported in Table 2. They indicate that state-of-the-art models beat the complexity-matched models by a margin of 12% accuracy. Therefore, for in-depth investigation of representations in scene-selective brain areas we used the top 3 models, i.e. HRNetv2, Upernet101, and Resnet101-PPM.

**Table 2:**
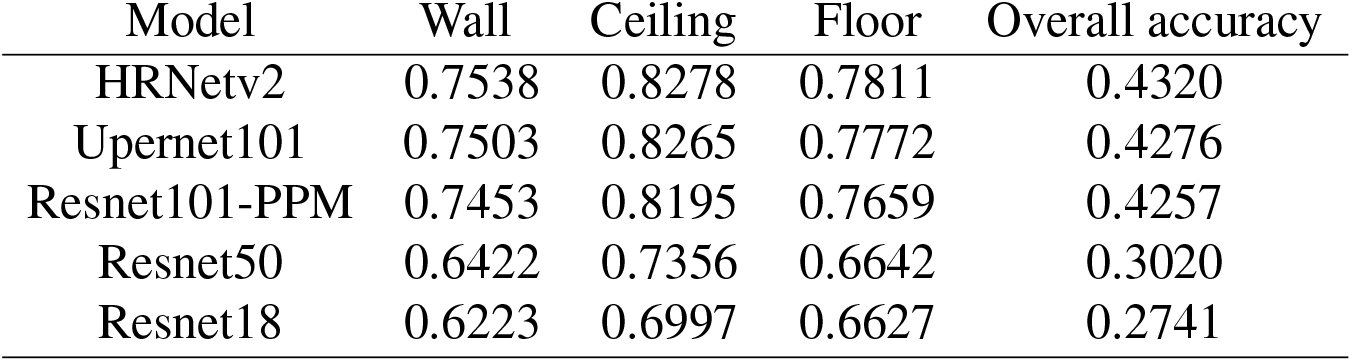
Scene parsing performance on ADE20k validation set. The tables shows the accuracy of detecting selected components along with overall accuracy for different scene parsing models in decreasing order.

### 3.3 Scene parsing networks reveal a differential contribution of wall, floor and ceiling components to representations in scene-selective regions

We investigated whether the scene components detected by a scene parsing DNN reveal a difference in the representational content of scene-selective ROIs. We focused on the three scene components - wall, floor, and ceiling - that were present in all the images of the stimulus set and compared them with scene-selective ROIs OPA and PPA.

We first report the RSA results (Figure 5a) of comparing a scene component RDM with OPA and PPA for three state-of-the-art architectures HRNetv2, Upernet101, and Resnet101-PPM. We found that the correlation of the OPA RDM with the floor RDM was significantly higher than of the wall and ceiling RDMs, and the correlation of the PPA RDM with the wall and floor RDMs was significantly higher than with the ceiling RDM. The above results held consistently across all investigated models. Together, this suggest that OPA and PPA have differential representational content with respect to scene components.

**Figure 5:**
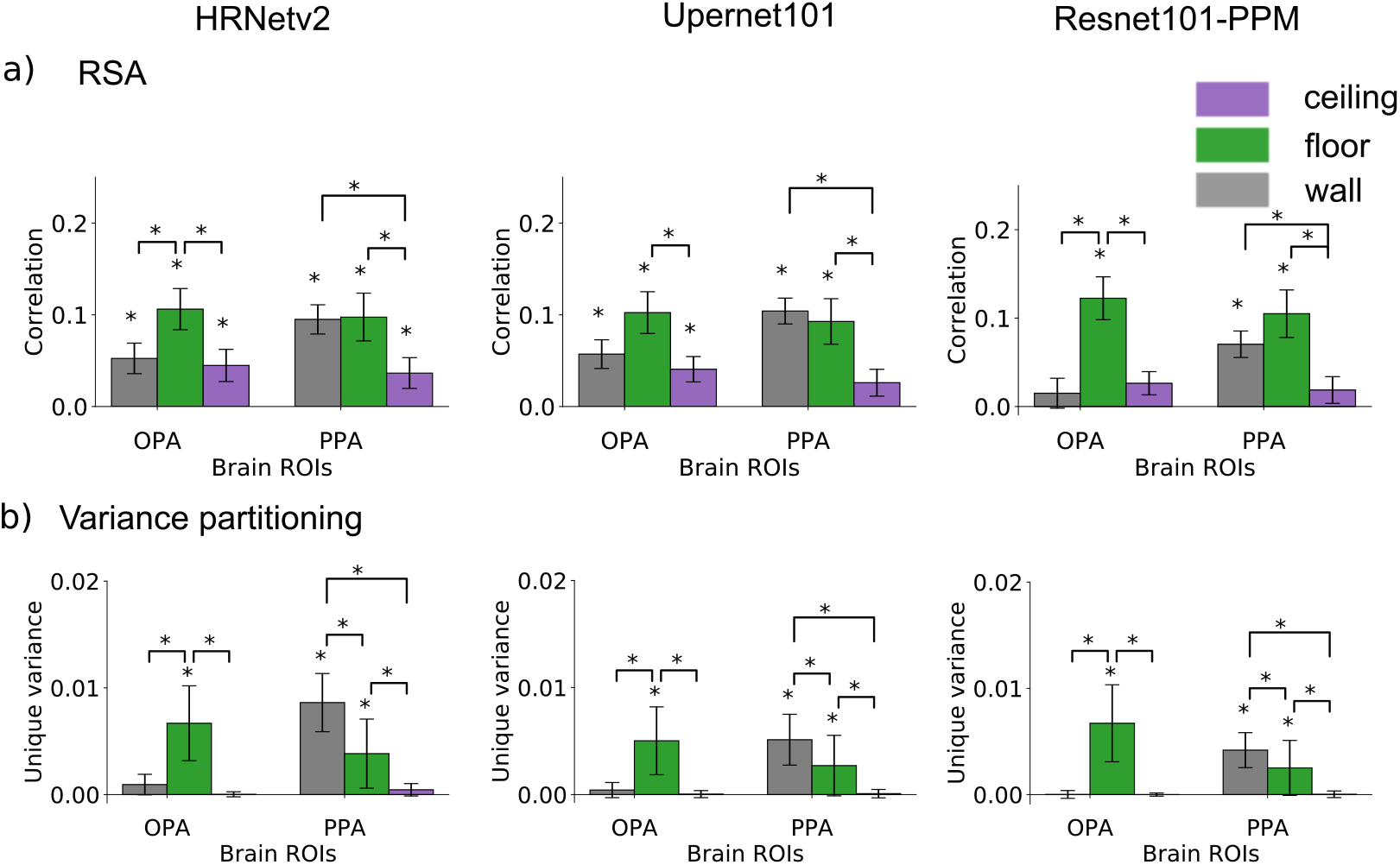
Scene components reveal the differences in representational content of OPA and PPA. a) Results of RSA between OPA and PPA with scene components of wall floor and ceiling for 3 state of the art models on scene parsing task HRnetv2 (left), Upernet101 (middle), Resnet101 (right), b) Unique variance accounted for in OPA and PPA by using components from HRnetv2 (left), Upernet101 (middle), Resnet101 (right) models. The asterisk at the top indicates the significance (*p <0.05) calculated by permuting the conditions 5000 times, FDR corrected with threshold equal to 0.05.

To tease out how much variance in OPA and PPA is explained by individual scene components, we apply variance partitioning to find the unique and shared variance of OPA and PPA RDMs explained by different scene component RDMs. We report the variance partitioning results showing unique variance explained by each component in Figure 5b and Venn diagram illustrating both unique and shared variances in Figure 6. We observed that in the case of OPA, the floor RDM explains significantly higher variance of OPA RDM uniquely compared to wall and ceiling RDMs. For PPA, the wall RDM explains significantly higher variance of PPA RDM uniquely compared to the floor and ceiling RDMs. Consistent with the RSA results above, this result reinforces the differences between OPA and PPA in representation of scene components.

**Figure 6:**
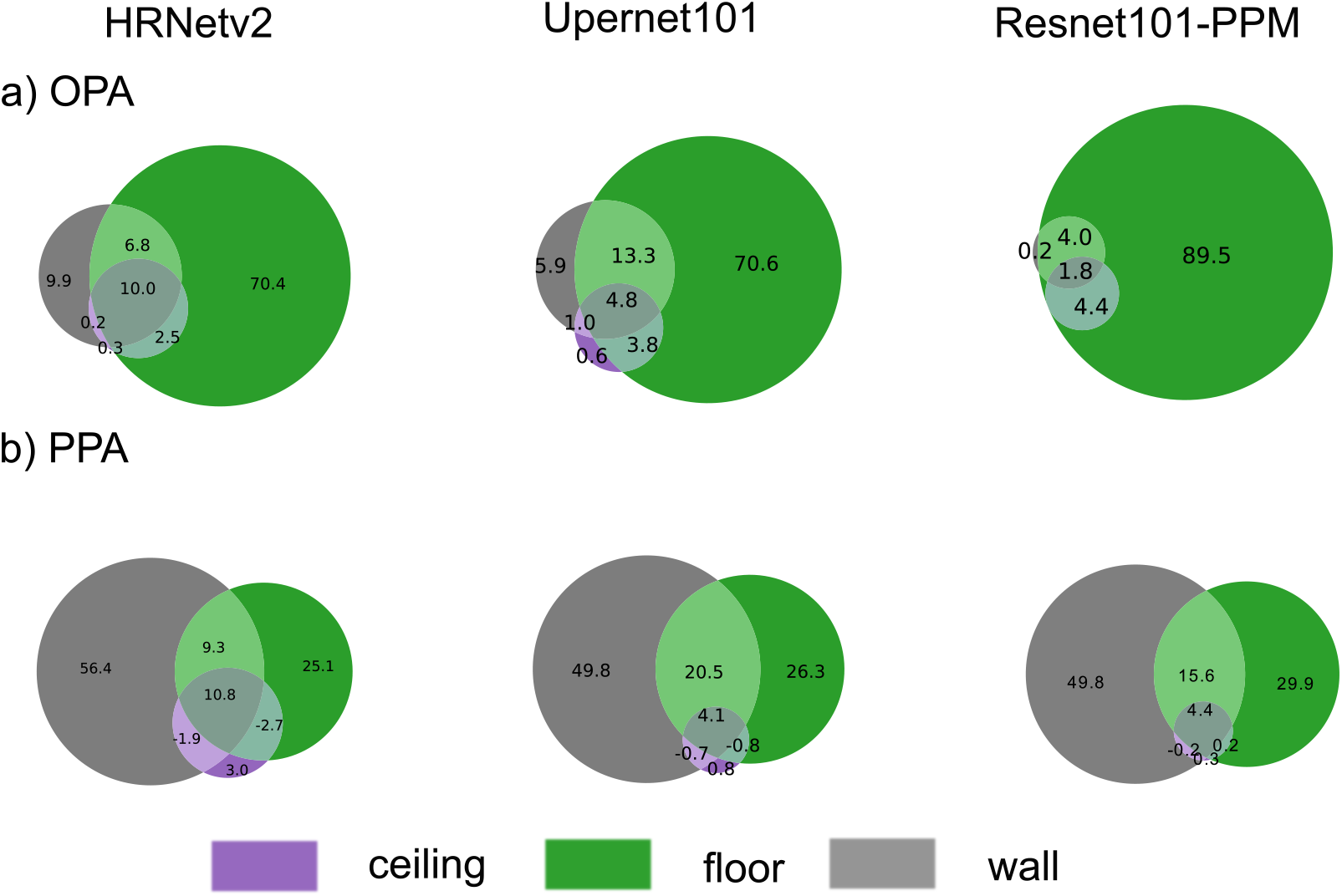
Variance Partition Results of Scene Components. The plots show the unique and shared variance of a) OPA and b) PPA explained by ceiling, wall and floor component for three architectures: HRNetv2 (left), Upernet101 (center), and Resnet101-PPM (right).

## 4 Discussion

In this study, we investigated the potential of scene parsing DNNs in predicting neural responses in scene-selective brain regions. We found that scene parsing DNNs predicted responses in scene-selective ROIs OPA and PPA better than scene-classification DNNs. We further showed that scene components detected by scene parsing DNNs revealed differences in representational content of OPA and PPA.

Previous work using DNNs to predict neural responses has emphasized the importance of the task for which the DNNs were optimized for [Yamins and DiCarlo, 2016, Khaligh-Razavi and Kriegeskorte, 2014, Richards et al., 2019]. We argue that the higher unique variance of scene-selective neural responses explained by scene parsing DNNs over scene-classification DNNs is due to such a difference in tasks. The scene classification task aims at identifying the category of the scene irrespective of the spatial organization of different components and objects in the scene. In contrast, the scene parsing task requires pixelwise labeling of the whole image and thus a more comprehensive understanding of the scene in terms of how different objects and components are spatially organized within a given scene. Our findings suggest that scene-selective ROIs OPA and PPA could be encoding additional information about the spatial organization of different components in the scene. This view is supported by evidence from neuroimaging literature [Kravitz et al., 2011, Park et al., 2011] showing that scene-selective regions represent the spatial layout of scenes.

Our in-depth analysis using scene components revealed differential representations in OPA and PPA. We observed that OPA had a significantly higher correlation with floor component as compared to ceiling and wall. One possible explanation for this high correlation with floor could be due to the sensitivity of OPA to stimulation in the lower visual field [Silson et al., 2015]. Another possible explanation for the observed difference could be due to OPA’s involvement in detecting navigational affordances [Bonner and Epstein, 2017]. Additionally, we found that PPA shows a significantly higher correlation with wall and floor compared to ceiling. The correlation of PPA with wall could possibly be attributed to its sensitivity to the upper visual field [Silson et al., 2015]. Since the ceiling was not represented, another plausible explanation could be that detecting wall is relevant to identifying the type of room, its texture [Henriksson et al., 2019, Park and Park, 2017] and landmarks for navigation [Troiani et al., 2012].

Previous work has already aimed at determining the nature of OPA representations by computational modelling [Bonner and Epstein, 2018]. For this, the authors determined for a DNN trained on scene categorization which individual DNN units most correlated with NAM and OPA and visualized those units using receptive field mapping and segmentation from Zhou et al. [2014]. The units extracted corresponded mostly to uninterrupted portions of floor and wall or the junctions between floor and wall. The results align with our findings that floor component explain OPA responses uniquely while the wall units could be attributed to shared variance explained by floor and wall components in our study. However, arguably segmentation extracted using the receptive field mapping method are less interpretable if we do not assign the unit to a particular scene component. Further, assigning a unit to a component based on receptive field mapping and segmentation maps is less accurate compared to the segmentation from a scene parsing DNN trained to segment components using ground truth segmentation maps. Thus, we believe due to highly accurate and interpretable segmentation maps, our approach to use scene-components generated using a scene parsing DNN is particularly well suited to reveal the representational content of scene-selective brain regions.

A limitation of our study is that the differences revealed between OPA and PPA is based on the analysis of only 3 scene components. This is due to the limitations of the stimulus set, which consistently had only 3 components that were present in all 50 images. Future work should exploit the full richness of scene components provided by DNNs trained on scene parsing. For this a stimulus set would have to be designed that contains many components in all images of the stimulus set. Another possible direction would be to use stimuli with annotations and use these annotations directly to compare with the fMRI responses.

To summarize, our findings provided evidence supporting the use of DNNs trained on the scene parsing task as a promising tool to predict and understand activity in the visual brain. We believe that this approach has the potential to be applied widely, providing interpretable results that give insights into how human visual cortex represents the visual world.

## Acknowledgements

We thank Agnessa Karapetian and Greta Häberle for their valuable comments on the manuscript. R.M.C. is supported by Deutsche Forschungsgemeinschaft (DFG) grants (CI241/1-1, CI241/3-1) and the European Research Council Starting Grant (ERC-2018-StG 803370). G.R. thanks the support of the Alfons and Gertrud Kassel Foundation.

